# Tumor-Infiltrating Lymphocytes Display Prognostic Signatures Associated with Chemotherapy Response in TNBC Patients

**DOI:** 10.1101/2024.11.01.621478

**Authors:** Shayantan Banerjee, Vijay K. Tiwari, Karthik Raman, Mohammad Inayatullah

**Author notes:** Present address: Centre for Digital Health, Indian Institute of Technology Bombay (Shayantan Banerjee).

## Abstract

Triple-negative breast cancer (TNBC) is an aggressive subtype often marked by resistance to neoadjuvant chemotherapy (NAC), making treatment particularly challenging. Tumor-infiltrating lymphocytes (TILs), crucial players in the immune landscape of tumors, have been associated with treatment outcomes, but the prognostic potential of TIL-derived gene markers in pre-NAC samples from TNBC patients remains understudied. In this research, we analyzed the single-cell transcriptional profiles of approximately 5,000 cells from four chemosensitive and four chemoresistant TNBC patients using publicly available datasets. Leveraging standard single-cell analysis, we identified differentially expressed gene signatures within the TIL subpopulation, highlighting significant immune activation pathways differentiating chemoresistant from chemosensitive tumors. By employing robust feature selection and repeated cross-validation across microarray and RNA-seq datasets, we developed a stable set of 30 TIL-based gene markers with notable prognostic relevance for NAC response in TNBC. These markers achieved an AUROC of 0.78 in the training set and validated with AUROCs of 0.8, 0.658, and 0.736 across five independent test datasets, demonstrating consistency across diverse platforms and sequencing technologies. Furthermore, increased expression of these gene signatures correlated with improved recurrence-free survival (RFS) in a cohort of 220 TNBC patients. This study enhances our understanding of the TIL transcriptional landscape in NAC response, identifying potential biomarkers and therapeutic targets for improving treatment outcomes in TNBC.

## Introduction

Breast cancer (BC) is one of the most prevalent cancers, affecting millions worldwide (1). The molecular classification of breast cancer subtypes plays a crucial role in diagnosis and treatment planning. This classification includes determining the expression of receptors such as estrogen (ER), progesterone (PR), and human epidermal growth factor receptor 2 (HER2) (2). Triple-negative breast cancer (TNBC), which accounts for approximately 15-20% of BC cases, is particularly aggressive due to its lack of ER, PR, and HER2 receptors. This subtype has a high risk of recurrence and poorer long-term survival rates (3). The ineffectiveness of endocrine and anti-HER2 therapies against TNBC often leaves chemotherapy and surgery as the primary treatment options (4). Neoadjuvant chemotherapy (NAC) offers a breast-conserving alternative by targeting the micro-metastatic components of the tumor. However, only about 50% of patients achieve pathological complete response (pCR) or recurrence-free survival (RFS) with this treatment. The remaining patients experience poor overall survival and a high risk of recurrence (5). Adopting a precision medicine approach, which includes understanding the TNBC transcriptome and identifying molecular markers, can significantly enhance patient stratification and clinical decision-making. By prioritizing patients’ responses to chemotherapy based on their molecular profiles, healthcare providers can improve treatment outcomes and tailor therapies to individual needs.

Several studies have explored the transcriptional landscape of TNBC at the bulk RNA-seq and single-cell level (2,6). Many studies have revealed that TNBC tumors harbor a high degree of heterogeneity due to cell differentiation, plasticity, cancer cell stemness, and tumor microenvironment (7,8). Kim et al. (9) studied the transcriptome of eight pre– and post-NAC-treated TNBC patients. They described two categories: “clonal extinction”, where tumor cells vanished after NAC treatment, and “clonal persistence,” where enrichment of tumor clones was observed post-treatment, indicating a lack of response. Tumor-infiltrating lymphocytes (TILs) are white blood cells that trigger an immune response after infiltrating the tumor cells. Recently, multiple studies have established an association between TIL abundance and the progression of TNBC (10–13). Specifically, a high abundance of TILs before treatment is usually associated with a high incidence of pathological complete response (pCR) (14). Similarly, at least 60% TILs in the residual disease after NAC treatment was associated with better outcomes in TNBC patients (15). Based on this data, we hypothesized that gene markers extracted from the TIL subpopulation of cells from single-cell expression measurements could be prognostic indicators for evaluating the NAC response in TNBC patients.

With the recent advancements in machine learning-assisted identification of gene biomarkers from publicly available expression datasets, its evaluation and use in patient stratification, therapy response predictions, and precision treatment design have increased (16–20). However, there is a strong unmet need to integrate single-cell and bulk RNA-seq expression measurements to identify predictors of response from the TIL subpopulation of cells belonging to TNBC patients. Therefore, in this study, we first extracted the cells belonging to the TIL subpopulation from single-cell data using established canonical markers and built machine-learning models to train and validate them on several independent test cohorts. Finally, the role of the established markers was explored using survival and pathway enrichment analysis.

## Materials and Methods

### Dataset selection and preprocessing

We analyzed a single-cell RNA sequencing (scRNA-seq) dataset comprising 5872 cells from matched pre– and post-treatment triple-negative breast cancer (TNBC) tumors (9). The dataset included samples from four patients who responded to neoadjuvant chemotherapy (NAC) with docetaxel and epirubicin (chemosensitive) and four patients who had residual disease (RD) after treatment (chemoresistant). The samples in this dataset were obtained from core biopsies taken before neoadjuvant chemotherapy (NAC) (0 cycles, pre-treatment), after two cycles of NAC, and from a surgical sample collected after four additional cycles of NAC combined with bevacizumab (post-treatment). Single-cell data was filtered to include only pre-treatment samples for further analysis. One particular cell with UMI KTN206_P_CCAAGCCATGG did not have corresponding phenotypic information available and was excluded from further analysis.

### Quality Control

The Seurat package (21) generated basic metadata for each cell in the count matrix. The PercentageFeatureSet() with the pattern MT^ was used to search for identifiers associated with the mitochondrial genes. The analysis was restricted to only protein-coding genes using the “HgProteinCodingGenes” gene list published by the Cell-ID package (22) in R. Following the methodology outlined in the original paper (9), we excluded cells that did not express either *GAPDH* or *ACTB* from the analysis. We also excluded a list of previously published 98 housekeeping genes (23). Natural log transformation with a pseudo count of one (i.e., ln(x+1)) of the already TPM-normalized data was performed, and the resulting expression matrix was used for the subsequent QC step. Steps outlined in the standard Seurat preprocessing workflow for generating metrics to include only high-quality cells for downstream processing were performed. Briefly, cells with unique feature counts >2500 or <200 or more than 5% mitochondrial content were excluded from the analysis. Similarly, genes with zero counts or those expressed in less than ten cells were removed. The clinical information about the patients stored in the metadata file of the filtered cell object included the chemotherapy labels (“chemosensitive” or “chemoresistant”) and the patient identity.

### Removing unwanted sources of variation, finding variable genes, and integrating single-cell data

The cleaned expression matrix was used to remove unwanted sources of variation that might cause cell clustering by artifacts in the downstream analysis. To understand whether cell cycle or mitochondrial content represents any significant source of variation, we chose the top 2000 highly variable genes, scaled the data, and performed principal component analysis. We then implemented two of the most common biological data correction measures by removing cell cycle effects and mitochondrial expression from the transcriptome using the ScaleData() with default parameters. Since we observed our cells clustering based on chemotherapy labels (“chemosensitive” vs. “chemoresistant”), we implemented the scRNA-seq integration workflow to remove the batch effect and align similar cells across different conditions using the highly variable genes (HVGs) (24). Briefly, we selected the most variable gene using the SelectIntegrationFeature(), prepared the Seurat object for the integration step with the PrepSCTIntegration(), canonical correlation analysis (CCA) with FindIntegrationAnchors() function, and final integration across chemotherapy labels with IntegrateData(). Uniform Manifold Approximation and Projection (UMAP) visualization was performed to compare the integrated and unintegrated expression matrix to check if the cells aligned well and if the dataset benefitted from the integration step.

### Clustering single-cell data, performing cell type annotation, and selecting tumor-infiltrating lymphocytes (TILs) subpopulation

The PCA scores derived from the expression measurements of 2000 highly variable genes from the integrated single-cell object were used to assign cells to clusters. We used the “Elbow method” to estimate the number of PCs to include in the clustering analysis. Cells were then clustered using the graph-based clustering method FindClusters(). To determine the granularity of the downstream clustering, “resolution” parameters 0.4, 0.5, 0.6, 0.8, 1.0, and 1.4 were provided as input to the function. The clustered cells were then projected into a 2D space and visualized using the RunUMAP() and the Dimplot().

Cell-type annotation was performed using the evidence-based scoring and annotation tool scCATCH (25) in R by setting the “species” parameter to “human” and the “tissue” parameter to a list containing “Breast,” “Mammary epithelium,” “Mammary gland,” “Blood,” and “Peripheral blood.” The findmarkergenes() and scCATCH() were employed to annotate each identified cluster automatically using a curated list of known cell markers in the tissue-specific reference database (CellMatch) (25). Identification of tumor-infiltrating lymphocytes was performed using established markers from the original paper (9). Marker genes CD8A, CD8B, and CD4 were used to represent the T cell subpopulation, and CD19 and MS4A1 were used to represent the B cell subpopulation. Cluster-wise distribution of TILs was denoted using expression data projected using VlnPlot() and FeaturePlot() in Seurat.

### Differential gene expression and functional enrichment analysis

Differentially upregulated marker genes with an absolute logFC cutoff of 1 and an adjusted *p*-value of 0.05 were identified using the FindAllMarkers() in Seurat. The ClusterProfiler (26) package in R was used to perform gene ontology (GO) analysis of the identified gene markers, and the corresponding Biological Process (BP), cellular components (CC), and molecular function (MF) terms were reported. Fifty hallmark genes were downloaded from the Molecular Signature Database (MSigDB) (27), and pathway scores were calculated for each cell using the AddModuleScore() in Seurat.

### Construction of the PPI network, identification of clusters and hub genes

The online STRING tool (version 12.0: https://string-db.org/) (28) analyzed the intersecting gene markers obtained from the integrated analysis of a differentially expressed and manually curated list of genes. A combined score cutoff of at least 0.4 was selected as the interaction confidence, and the results were visualized using the Cytoscape software (v. 3.20.2). MCODE (v.2.0.3) was used to identify highly interconnected modules from the network (MCODE score >= 5). The top 20 hub genes were selected using seven standard algorithms (MCC, Degree, Closeness, Radiality, Stress, and EPC) available in the Cytohubba plugin of Cytoscape, and the shared genes were considered for further analysis.

### Identification of potential marker genes using prior biological knowledge

Performing integrative analysis using already published gene lists with prior biological knowledge has been shown to improve the task of gene selection for biomarker detection (29). To identify a small set of relevant features for machine learning analysis, we manually curated a set of marker genes predictive of NAC response among breast cancer patients (Table 1). Some studies developed RNA classifiers using derivation and validation datasets derived exclusively from TNBC patients (30–34) or specific subtypes such as ER+/HER2-(35). The remaining studies included all subtypes based on ER/PR/HER2 status (19,36–40). Unique gene symbols pooled from the aforementioned studies were standardized using the HGNC Multi Symbol checker (https://www.genenames.org/tools/multi-symbol-checker/). In summary, a total of 579 marker genes related to NAC response were collected using this process.

**Table 1:**
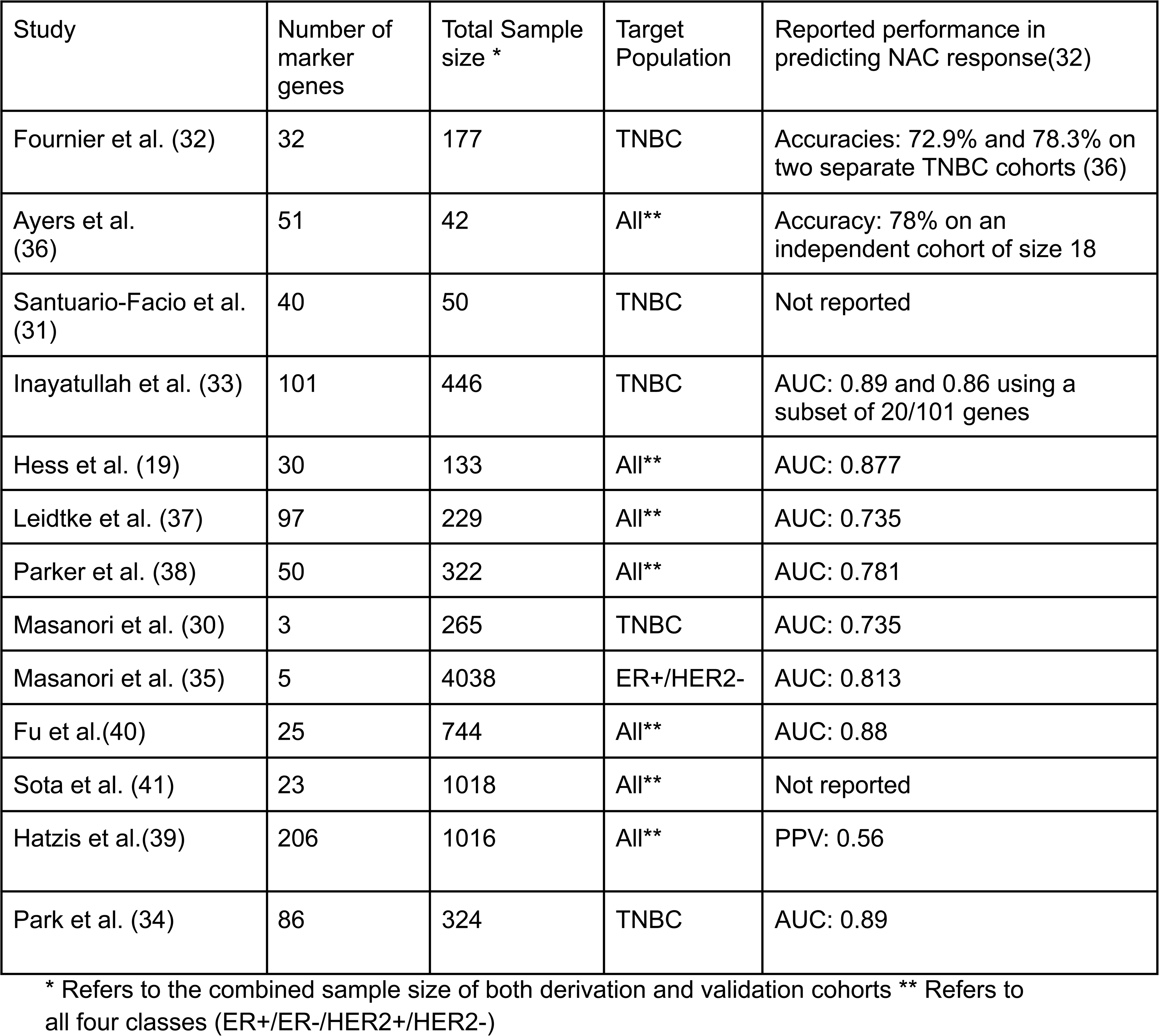
Summary of studies predicting NAC response using gene markers.

| Study | Number of marker genes | Total Sample size * | Target Population | Reported performance in predicting NAC response(32) |
| --- | --- | --- | --- | --- |
| Fournier et al. (32) | 32 | 177 | TNBC | Accuracies: 72.9% and 78.3% on two separate TNBC cohorts (36) |
| Ayers et al. (36) | 51 | 42 | All** | Accuracy: 78% on an independent cohort of size 18 |
| Santuario-Facio et al. (31) | 40 | 50 | TNBC | Not reported |
| Inayatullah et al. (33) | 101 | 446 | TNBC | AUC: 0.89 and 0.86 using a subset of 20/101 genes |
| Hess et al. (19) | 30 | 133 | All** | AUC: 0.877 |
| Leidtke et al. (37) | 97 | 229 | All** | AUC: 0.735 |
| Parker et al. (38) | 50 | 322 | All** | AUC: 0.781 |
| Masanori et al. (30) | 3 | 265 | TNBC | AUC: 0.735 |
| Masanori et al. (35) | 5 | 4038 | ER+/HER2- | AUC: 0.813 |
| Fu et al.(40) | 25 | 744 | All** | AUC: 0.88 |
| Sota et al. (41) | 23 | 1018 | All** | Not reported |
| Hatzis et al.(39) | 206 | 1016 | All** | PPV: 0.56 |
| Park et al. (34) | 86 | 324 | TNBC | AUC: 0.89 |

\* Refers to the combined sample size of both derivation and validation cohorts \*\* Refers to all four classes (ER+/ER-/HER2+/HER2-)

### Bulk transcriptome data analysis for model building and validation

We selected five microarray datasets of pre-treated TNBC patients with NAC response labels to train and test our RNA-based classifiers from the GEO database. Table 2 summarizes the characteristics of the derivation and validation sets, including the number of patients, sequencing platform, and NAC regimen used in this study. Owing to the large number of TNBC samples in the GSE25066 dataset, we considered this the derivation dataset and the remaining four validation datasets. Due to the absence of response labels, we considered 3-year event-free survival (EFS) as the surrogate predictor variable for the dataset GSE31519. Patients exhibiting complete EFS were considered to be in a pathological complete response state and vice versa. The ReadAffy() from the “Affy” package (42) was used to load the raw expression values in R, and the “gcrma” package (43) was used to perform the Robust Multi-Average (RMA) parameter method of normalization and background correction. For the microarray datasets, GSE25066, GSE20271, GSE20194, and GSE31519, the Affymetrix probes were matched to gene symbols using the Affymetrix Human Genome U133 Plus 2.0 (hgu133plus2.db) and for GSE106977, the Affymetrix Human Transcriptome Array 2.0 was used to derive the gene symbols. The average expression values of multiple probes that match the same gene were considered. The three validation datasets of the GPL96 platform were combined into one large validation set. The resulting batch effect was removed using the Combat() from the “sva” package (44) in R. Using the expression values, the before and after boxplots and scatter plots with the first two principal components were constructed to show the effect of normalization and batch effect correction, respectively. The normalized TPM count matrix of the RNA-seq dataset GSE163882 was downloaded from the GEO database. We used the gene annotations deposited by the authors and subsetted only TNBC samples to prepare the processed data matrix for further analysis.

**Table 2:** Description of the derivation and validation datasets used in this study.

| Category | GEO accession | #TNBC samples | Prediction labels | Data distribution | Platform ID | NAC regimen | Clinical Variables available |
| --- | --- | --- | --- | --- | --- | --- | --- |
| Derivation (microarray) | GSE25066 (39) | 170 | pCR/RD | pCR: 57<br>RD: 113 | GPL96 | TFAC and ACT | Age, Nodal Status, Clinical stage |
| Validation (microarray) | GSE20271 (17) | 59 | pCR/RD | pCR: 13<br>RD: 46 | GPL96 | PFCD | Age, Tumor Grade |
| Validation (microarray) | GSE20194 (45) | 71 | pCR/RD | pCR: 25<br>RD: 46 | GPL96 | PFCD | Age, Tumor Grade |
| Validation (microarray) | GSE31519 (46) | 64 | 3-year Event-Free Survival (EFS) | Yes: 36<br>No: 28 | GPL96 | Not Applicable | Tumor Grade |
| Validation (microarray) | GSE106977 (47) | 119 | pCR/RD | pCR: 43<br>RD: 76 | GPL17586 | A/T or A/T+CA | Treatment information |
| Validation (RNA-seq) | GSE163882 (48) | 90 | pCR/RD | pCR: 32<br>RD: 58 | GPL18573 | A/T | Age, Tumor Grade, Stage |

### Feature selection and training

The processed microarray derivation dataset had 13,287 genes for 170 samples. The first step of feature reduction included selecting common genes between the list of genes differentially expressed among the TIL subpopulation derived from the single-cell data and the 579 NAC-related marker genes based on existing literature. This resulted in a reduced list of 226 genes for further analysis. Our next objective was to reduce the dataset’s dimensionality by removing redundant variables. Owing to the imbalance in the dataset, we performed a 5-fold stratified cross-validation by randomly partitioning the dataset into five equal parts, ensuring that the proportion of labels is intact in each train-test split. During every run of the cross-validation experiment, we employed recursive feature elimination (RFE) using three estimators: random forests, linear SVM, and logistic regression, resulting in three feature sets: RF-RFE, SVM-RFE, and LOGIT-RFE. This process is repeated 100 times, resulting in 500 unique train-test splits, ensuring a different dataset split during every run. The fraction of times a particular gene was selected out of 500 runs was used to rank the genes in decreasing order of strength of association with the predictor variable. The Borda count ranking aggregation method combined the three ranked lists (RF-RFE, SVM-RFE, and LOGIT-RFE) and generated a single list of highly discriminative features ranked in decreasing order of relevance. The top ‘n’ genes (n = 10,15,20,25,30), also known as stable genes, were used to train and tune the machine learning models.

### Sampling techniques, Model selection, and Tuning

Datasets with a skewed distribution of labels can negatively affect machine learning algorithms. This study used various resampling techniques to balance the derivation dataset (Supplementary Table S1). Hyperparameter tuning of six binary classifiers was implemented using a 5-fold stratified cross-validated grid search technique repeated 20 times over a parameter grid using the Mathews correlation coefficient (MCC) metric as the scoring function. The top-performing model was extracted using the best_estimator_ argument in scikit-learn and stored for subsequent analysis. The entire grid of parameters and classifier settings is available in Supplementary Table S1B. Due to the imbalance, we experimented with classification thresholds between 0 and 1 with step sizes of 0.001. The model performances at the threshold with the maximum MCC were reported. The tuning step utilized the derivation dataset to find the best feature set (n=10,15…30), sampling technique, and classifier combination. The top-trained models were then directly applied to the validation datasets to measure the generalizability of the classifiers. We assessed the performance of our classifiers using four widely used performance metrics: Sensitivity, Specificity, Mathews correlation coefficient, Area under the precision-recall curve (AUPRC), Area under the ROC curve (AUROC), and F1-score.

### Survival analysis

We performed survival analysis of the top gene predictors using the online KMPlotter tool (https://kmplot.com/analysis/index.php?p=service). Using the gene symbols in the “input multiple genes” box and selecting “mean expression of genes” (to ensure a single plot for all 30 genes), we selected ER, PR (IHC), and HER2 array status as negative. We selected RFS as the clinical endpoint. This led to a subset of 220 TNBC patients with survival information and expression values.

### Machine learning analysis – Validation

Both the derivation and unseen independent validation datasets were first scaled between 0 and 1 using the min-max scaling technique. However, we only used the training parameters to scale the test set. We use fit_transform() from scikit-learn on the training data but only the transform() on the test set. Next, the top-tuned model derived earlier, consisting of the optimal number of ‘n’ stable genes, sampling-classifier technique, and classification threshold, was used to generate predictions on the four validation sets. The ROC curve and 95% confidence intervals for the classification metrics were obtained through 1000 iterations using repeated sampling with a replacement approach.

### Statistical Analysis

A Wilcoxon rank sum test was used to find differentially expressed genes using the FindAllMarrkers() in Seurat. Correction for multiple hypothesis testing was performed using the Benjamini Hochberg (BH) technique. A Mann-Whitney U test (p-value cutoff of 0.05) was performed to find the statistical significance between the expression levels of different genes.

## Results

### Single-cell analysis revealed transcriptional signatures associated with immune cell activation within chemosensitive TNBC tumors

To identify transcriptional signatures of prognostic value, we utilized single-cell transcriptome datasets of 2640 pre-treated cells from “chemoresistant” (n=4) and “chemosensitive” (n=4) (Figure 1A) TNBC patients. Standard preprocessing steps outlined in the Seurat workflow (21) were applied to the raw single-cell count data. We aimed to develop a filtering mechanism to retain high-quality cells for feature selection. The “Quality Control” steps outlined in the “Methods” section fetched a final processed dataset of 14,330 genes and 2034 cells. The effect of single-cell data integration was visualized using UMAP plots based on NAC response labels and patient identities (Supplementary Figure 1A-D). Unsupervised clustering using optimal resolution parameters of 964 chemoresistant and 1070 chemosensitive pre-treated single cells identified nine unique clusters (Figure 1B; Supplementary Table S2). Automatic cell type annotation using scCATCH identified cancer stem cells, regulatory T cells, Plasmacytoid dendritic cells, naive T cells, luminal epithelial cells, and mammary epithelial cells (Figure 1C).

**Figure 1:**
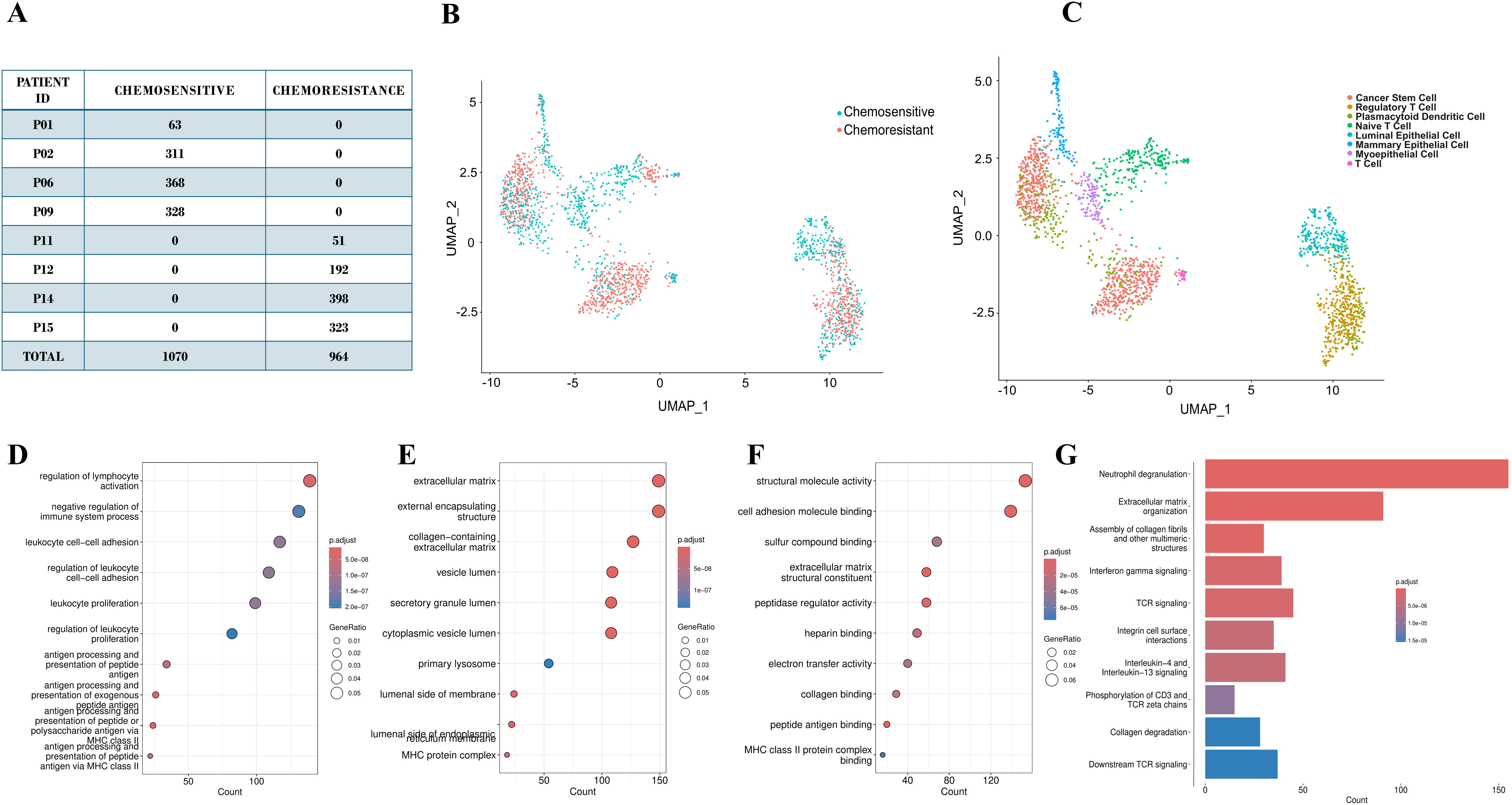
Results from the single-cell analysis using the raw count data from 2640 pre-treated cells belonging to 8 patients (4 chemosensitive and 4 chemoresistant). (A) Distribution of cells across the two patient groups. (B) UMAP plot of the integrated dataset consisting of 2640 cells (C) Single-cell type annotation using the scCATCH tool. (D) Gene ontology analysis using the differentially expressed genes across all 2640 cells showing terms enriched in terms of biological processes (BP) (E) Cellular components (CC) and (F) Molecular functions (G) Significant REACTOME pathways altered using the same set of DEGs identified across 2640 cells.

To derive a generalizable gene signature differentiating between chemoresistant and chemosensitive patients, first, we compared the transcriptome of the two patient groups and identified differentially expressed genes (Supplementary Figure 2A). Gene Ontology analysis revealed regulation of lymphocyte activation (GO:0051251), negative regulation of the immune system process (GO:0002683), and leukocyte cell-cell adhesion (GO:0007159) as some of the significantly enriched biological processes for genes that were upregulated in chemoresistant vs. chemosensitive cells (Figure 1D-F). These proteins were primarily located on the extracellular matrix (GO:0031012) and vesicle lumen (GO:0031983), as indicated by the significantly altered cellular components and structural molecular activity (GO:0005198) and cell adhesion molecule binding (GO:0050839) as some of the significantly enriched molecular functions affected (Figure 1D-F). Neutrophil degranulation (R-HSA-6798695), interferon-gamma signaling (R-HSA-877300), and Interleukin-4 and Interleukin-13 signaling (R-HSA-6785807) were some of the significantly altered REACTOME pathways (Figure 1G; Supplementary Table S3A). These findings suggest that immune-related pathways and cell adhesion mechanisms play a critical role in driving the chemoresistant phenotype in TNBC, providing potential targets for therapeutic intervention.

### Treatment naïve tumor-infiltrating lymphocyte (TIL) subpopulation exhibits distinct transcriptional programs in chemoresistant and chemosensitive tumors

To identify whether expression differences in cell subpopulations derived from single-cell data can be used to differentiate between chemosensitive and chemoresistant samples, we extracted tumor-infiltrating lymphocytes (TILs) from 2034 pre-treated cells. The UMAP plots (Figure 2A) show the expression levels of marker genes CD8A, CD8B, and CD4 for T cells and CD19 and MS4A1 for B cells across the identified tumor-infiltrating lymphocytes (TILs) clusters. The cluster-wise distribution indicates distinct regions of T and B cell subpopulations, highlighting the heterogeneity of immune cell infiltration in the tumor microenvironment. A total of 806 cells from the TIL population were present in all nine clusters identified in the previous section. The volcano plot (Figure 2B) highlights the differentially expressed genes (adjusted p-val < 0.05; absolute log fold change cutoff of 1; Supplementary Figure 2B; Supplementary Table S4), including HLA-DRA and STAT3, in the TIL population. Gene ontology analysis of the differentially expressed upregulated genes identified regulation of lymphocyte activation (GO:0051251), negative regulation of immune system process (GO:0002683), and leukocyte cell-cell adhesion (GO:0007159) as the top enriched biological processes (Figure 2C). This is in line with earlier reports indicating the presence of an active antitumor immune response in TNBC patients mediated by high expression of TIL-based marker genes (49,50). These markers play a crucial role in modulating the effect of NAC on the tumor microenvironment.

**Figure 2:**
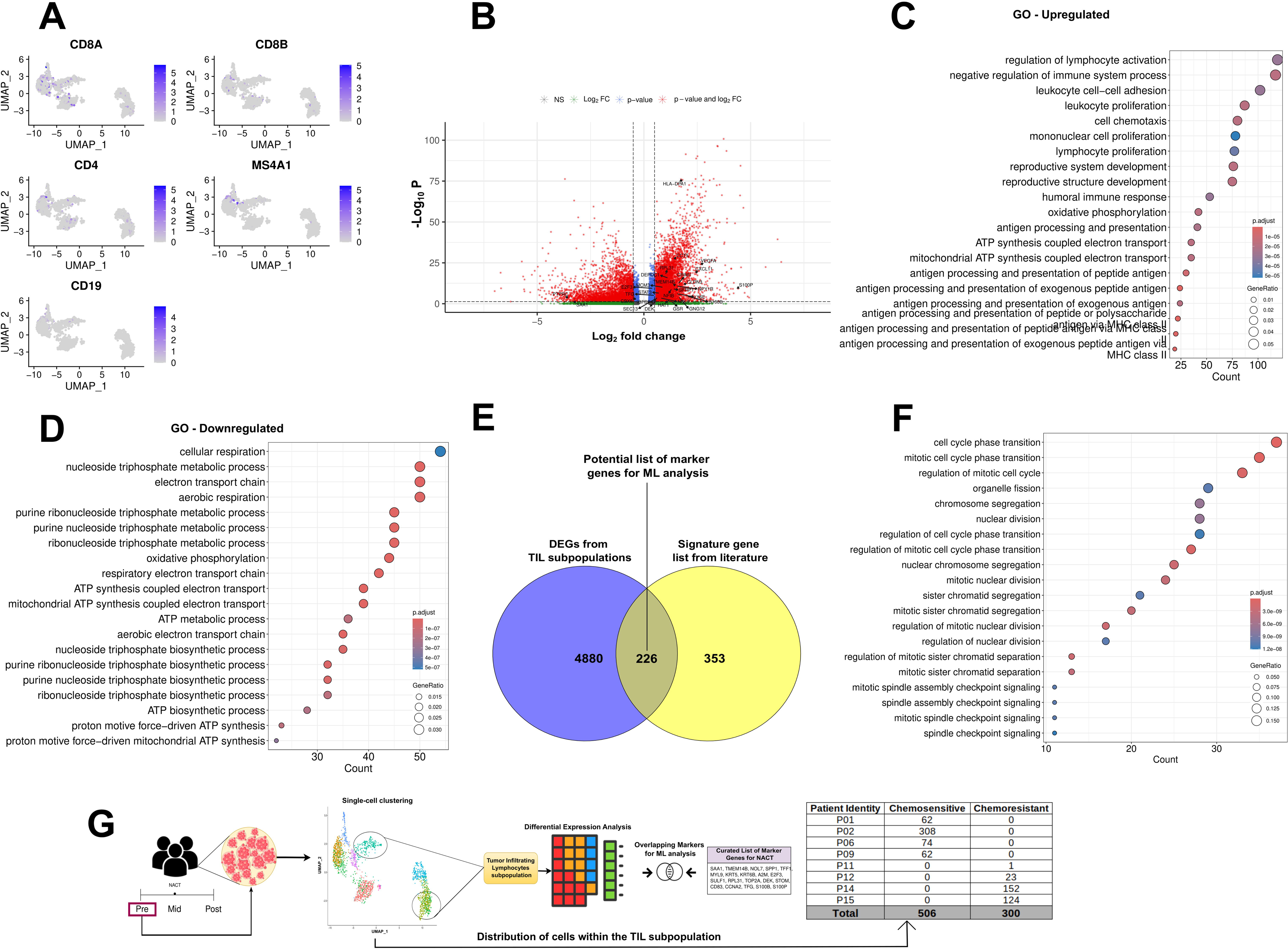
Results from the single-cell analysis restricted to 806 cells from the TIL subpopulation. (A) Feature plot showing the distribution of TILs across all nine single-cell clusters (B) Volcano plot highlighting the differentially expressed genes (adjusted p-val < 0.05; absolute log fold change cutoff of 1) within the TIL subpopulation (C) Gene ontology analysis using the same upregulated and (D) downregulated genes. (E) Venn diagram showing the intersection between the two groups of genes – differentially expressed in the TIL subpopulation and signature gene list from the literature. (F) Gene ontology analysis results using the top 226 markers (G) Workflow outlining the identification of tumor-infiltrating lymphocyte (TIL) subpopulations, highlighting the differential cell distribution between chemosensitive (506 cells) and chemoresistant (300 cells) patients, with a curated list of overlapping marker genes selected for further machine learning analysis

Similarly, cellular respiration (GO:0045333), nucleoside triphosphate metabolic process (GO:0009141), and electron transport chain (GO:0022900) were the enriched GO terms for the downregulated genes (Figure 2D). This observed downregulation of metabolic pathways may reflect an overall impaired metabolic capacity of TILs in chemoresistant tumors due to metabolic reprogramming or exhaustion, potentially contributing to their reduced antitumor activity in TNBC patients (51). Some of the altered pathways from the REACTOME database (Supplementary Table S3B) included Cytokine Signaling in the Immune system (R-HSA-1280215), Neutrophil degranulation (R-HSA-6798695), Interferon alpha/beta signaling (R-HSA-909733) and Programmed cell death (R-HSA-5357801). These alterations highlight the complex interactions between TILs and the tumor microenvironment in TNBC and underscore how tumors can modulate immune cell function to influence response to therapy.

The workflow in Figure 2G outlines the identification of tumor-infiltrating lymphocyte (TIL) subpopulations, highlighting the differential cell distribution between chemosensitive (506 cells) and chemoresistant (300 cells) patients, with a curated list of overlapping marker genes selected for further machine learning analysis. The intersection between differentially expressed genes from pre-treated cells belonging to the TIL subpopulation and a curated set of marker genes (Supplementary Table S5A) predictive of NAC response among breast cancer patients (Table 1) derived 226 potential markers (Supplementary Table S5B) and led to the reduction of the feature space for ML analysis (Figure 2E). The PPI network for the 226 potential marker genes was constructed using the STRING tool and exported to Cytoscape. GO enrichment revealed terms associated with cellular division, including chromosome segregation (GO:0007059), nuclear division (GO:0000280), and organelle fission (GO:0048285) (Figure 2F).

### Network Analysis of Shared TIL and NAC Response Gene Signatures Highlights Cell Cycle Regulation as a Key Enriched Pathway

Network analysis of shared gene signatures is essential for identifying key pathways in complex biological processes. This approach provides a deeper understanding of how different gene sets contribute to specific cellular responses, ultimately aiding biomarker discovery and therapeutic development. The MCODE plugin of the Cytoscape tool identified significant modules of the PPI network—one module with an MCODE score of 41.57 consisted of 43 nodes and 873 edges (Figure 3A; Supplementary Table S6A). GO analysis showed that the proteins in this module were related to cell cycle regulation, microtubule binding, and protein binding. Using the Cytohubba plugin, the top 20 hub genes were calculated using seven different algorithms, and their overlaps were assessed using an Upset plot (Supplementary Table S6B; Figure 3B(i-vii), 3C). Five common hub genes, selected by six out of seven algorithms, included forkhead box M1 (FOXM1), cyclin B1 (CCNB1), cyclin A2 (CCNA2), baculoviral IAP repeat containing 5 (BIRC5), and DNA topoisomerase II alpha (TOP2A). Table 3 shows their full names and related functions. Survival analysis of the hub genes on a subset of TNBC patients using the KMplotter tool revealed a significantly poorer survival linked to high expression of all five hub genes (Figure 3D; Table 3).

**Figure 3:**
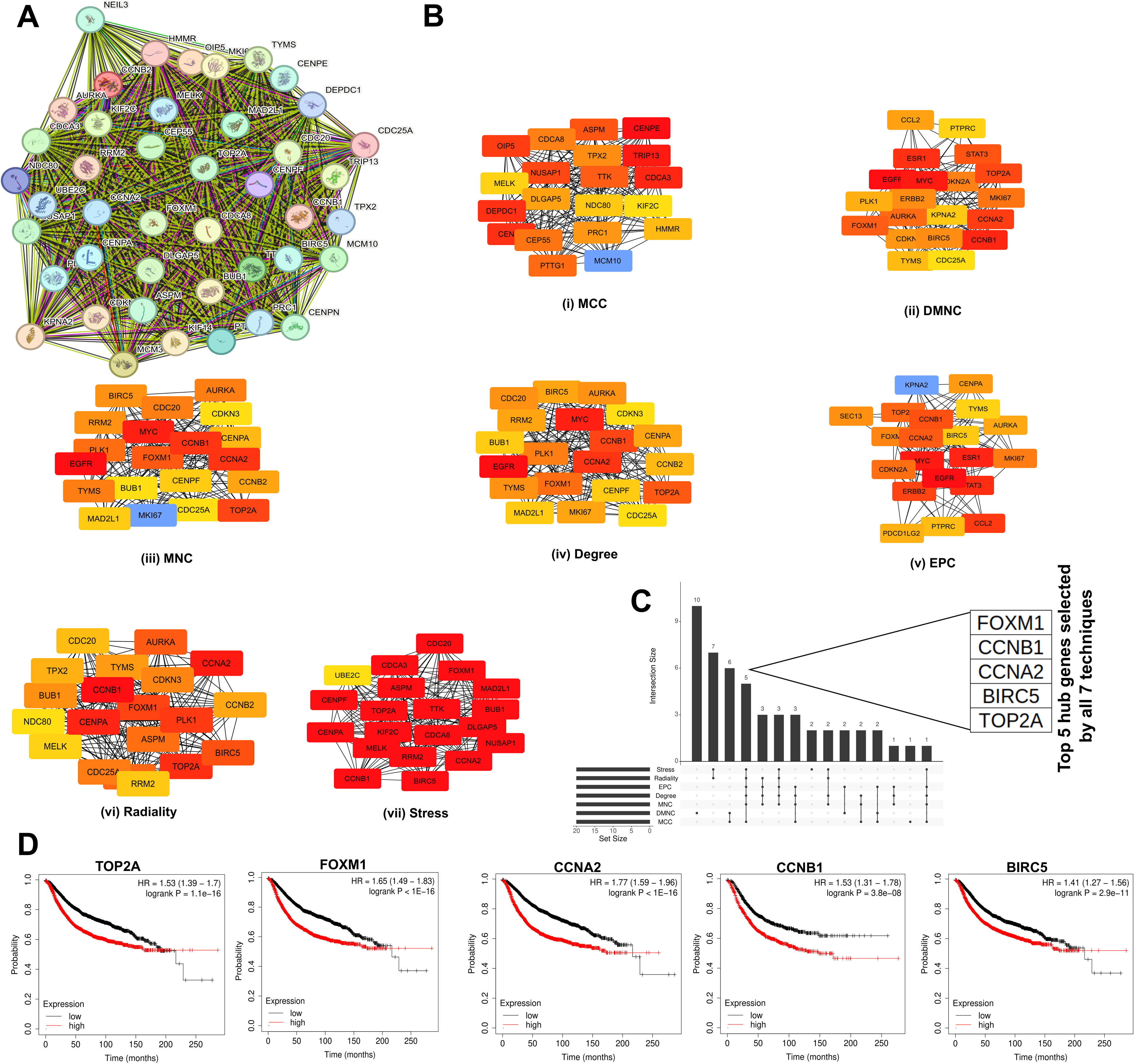
Network Analysis of Shared TIL and NAC Response Gene Signatures (A) PPI network for the 226 potential marker genes was constructed using the STRING tool and exported to Cytoscape. (B) (i-vii) The top 20 hub genes were derived using the Cytohubba plugin, and consensus was generated using seven different algorithms (C) The overlap between the seven algorithms was assessed using an Upset plot. (D) Survival analysis of the common hub genes and their association between expression and survival was visualized using the KMPlotter tool.

**Table 3:** List of hub genes and their names and functionalities.

| Gene Symbol | Gene Name | Function | Survival analysis |  |
| --- | --- | --- | --- | --- |
|  |  |  | Hazard Ratio | Log-Rank P value |
| FOXM1 | Forkhead box M1 | The protein encoded by this gene is a transcriptional activator involved in cell proliferation. | 1.65<br>(1.49-1.83) | <1e-16 |
| CCNB1 | Cyclin B1 | The protein encoded by this gene is a regulatory protein involved in mitosis. | 1.52<br>(1.31-1.78) | 3.8e-08 |
| CCNA2 | Cyclin A2 | The protein encoded by this gene belongs to the highly conserved cyclin family, whose members function as cell cycle regulators. | 1.77<br>(1.59-1.96) | <1e-16 |
| TOP2A | DNA topoisomerase II alpha | This gene encodes a DNA topoisomerase, an enzyme that controls and alters the topologic states of DNA during transcription. This nuclear enzyme is involved in chromosome condensation, chromatid separation, and the relief of torsional stress that occurs during DNA transcription and replication. | 1.53<br>(1.39-1.7) | 1.1e-16 |
| BIRC5 | Baculoviral IAP repeat containing 5 | This gene is a member of the inhibitor of apoptosis (IAP) gene family, which encodes negative regulatory proteins that prevent apoptotic cell death. | 1.41<br>(1.27-1.56) | 2.9e-11 |

### Robust Feature Selection followed by Subsequent Validation of Machine Learning Classifiers Identify TIL-Associated Gene Markers as Prognostic Indicators of NAC Response with High Predictive Performance

Machine learning algorithms facilitate biomarker discovery by identifying complex nonlinear associations between genes and the phenotype of interest from high-dimensional expression measurements. Owing to the lack of single-cell datasets with annotated data on pre-treated NAC-administered TNBC patients, we employed recently published microarray and RNA-sequencing datasets to train and validate our list of markers. Figure 4A shows the overall workflow of the machine learning analysis. The three major components involved the 1) Feature extraction phase, where we identified a ranked list of features from a single microarray dataset; 2) Model building phase, where we built classification models employing repeated stratified 5-fold cross-validation using different combinations of classifiers and the number of top-ranked features; and the 3) Model validation phase: in which we applied the classification models derived from step 2 on unseen, independent test sets to estimate the generalizability of the identified markers.

**Figure 4:**
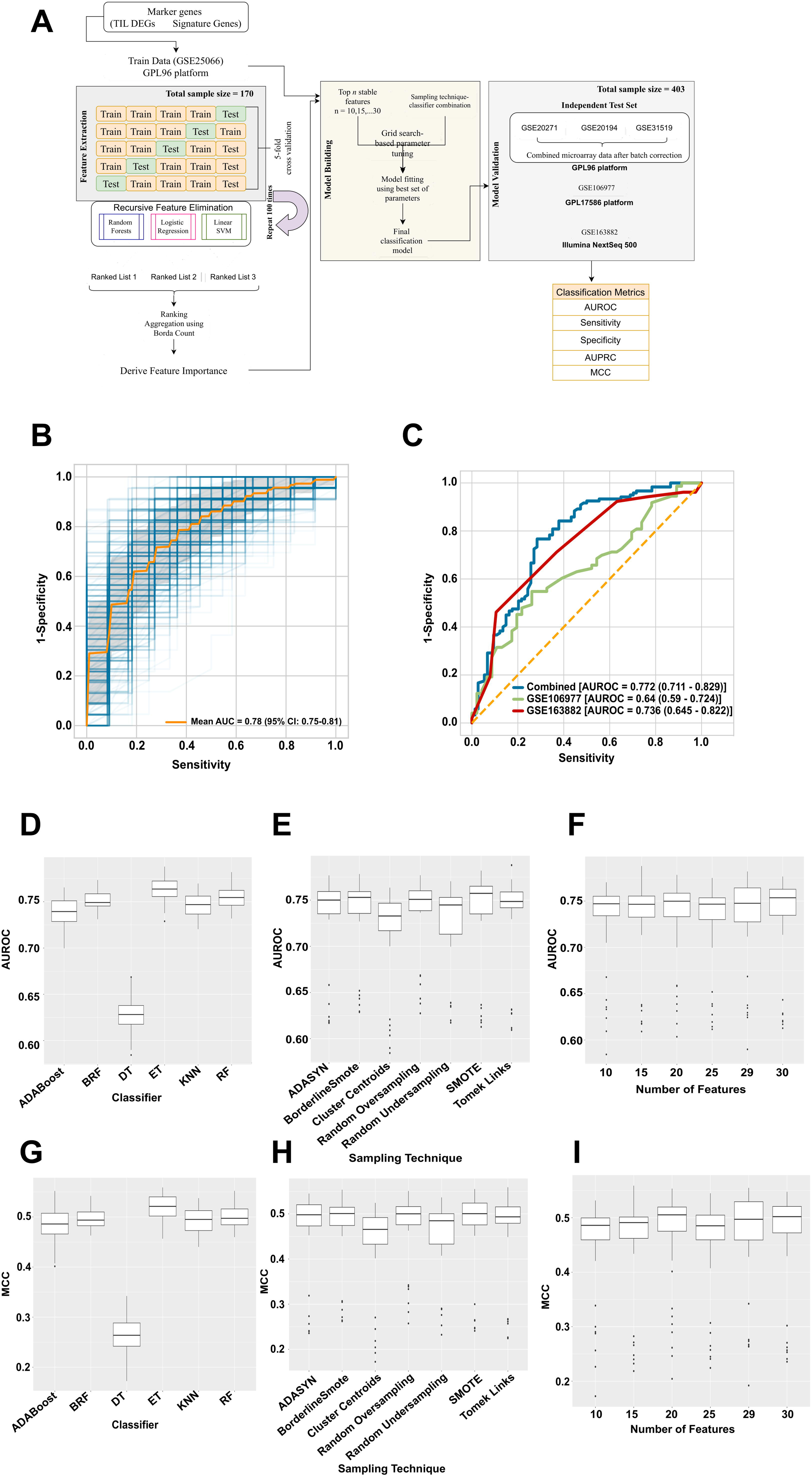
Machine learning derivation and validation of marker genes. (A) Overall machine learning workflow to derive a set of generalizable features using microarray dataset and validating the same using independent microarray and RNA-seq datasets. (B) Internal Validation area under the ROC curve (C) External validation area under the ROC curve (D-F) The variation in the performance, in terms of AUC for different classifiers, sampling techniques, and top-ranked features. (G-I) The variation in the performance, in terms of MCC for different classifiers, sampling techniques, and top-ranked features.

The first step of the machine learning analysis involved deriving a ranked list of features for further model building and validation. We performed repeated cross-validation experiments and employed robust rank aggregation techniques to identify a subset of consistently selected 30 stable genes sorted based on importance for training and validation (Supplementary Table S5C). Using the top *n* (n=10,15,20…30) stable genes (NOL7, HAT1, VEGFA, NFIB, S100B, DEPDC1, STAT3, S100P, NPY1R, GSR, SEC13, TFG, CCT5, CRIM1, SFRP1, GNG12, TMEM14B, PTPRC, CDKN2A, E2F3, IGFBP4, CLDN3, RPL31, MCM3, SAA1, CXCL11, LY6E, DEK, HLA-DPA1, CBX6), we trained different binary classifiers using stratified 5-fold cross-validation and derived the corresponding binary classification metrics. Within each run of the cross-validation experiments, we experimented with six classifiers, seven sampling techniques, and six feature sets, leading to a total of 252 combinations. We repeated the 5-fold cross-validation process 20 times, leading to 100 unique train-test splits. The variation in the performance, in terms of AUC and MCC for different classifiers, sampling techniques, and top-ranked features, is shown in Figure 4D-I. Extra trees classifier, random forests, and k-nearest neighbors were the top three classifiers with MCCs 0.52 (0.51-0.54), 0.497 (0.489-0.509) and 0.495 (0.479-0.501), respectively (Figure 4G). The difference in the MCC was significant (Wilcox test; p < 0.05) between extra trees and random forests and extra trees and k-nearest neighbors but not significant between random forests and k-nearest neighbors. Similarly, BorderlineSMOTE, SMOTE, and random oversampling achieved an overall MCC of 0.5 (0.48-0.51). However, no significant (Wilcox test; p < 0.05) difference in MCC was observed between the three techniques (Figure 4H). Finally, the top 20, 29, and 30 features achieved the highest MCC of 0.509 (0.488-0.511), 0.502 (0.484-0.515) and 0.497 (0.481-0.515), respectively. No significant differences were observed in terms of MCC.

We further validated the internal validation performance of the top sampling-classifier-feature set combination using several microarray-based independent expression datasets. The detailed results for all the combinations can be found in Supplementary Table S7. In the combined dataset (GSE20271, GSE20194, and GSE31519) and GSE106977, the best results were obtained using the top-performing combination from the internal validation results (Table 4A), i.e., random oversampling, extra trees, and top 30 ranked features which belonged to both TNBC-specific and non specific gene lists (Supplementary Table S5C). The corresponding classification metrics such as sensitivity, specificity, AUPRC, AUROC, F1 score, and MCC were 0.908, 0.527, 0.744, 0.772, 0.751, 0.483 and 0.534, 0.739, 0.694, 0.64, 0.616, 0.269, respectively (Table 4B). Owing to feature-wise distributional differences between the two sequencing technologies (microarray and RNA-seq), we observed the top performances for the RNA-seq dataset (GSE163882) using a different combination – *K*-nearest neighbors, Tomek Links, and 29 top features (Supplementary Table S8). This model was still one of the top 10 models reported from the internal validation analysis and resulted in a sensitivity of 0.923, specificity of 0.368, AUPRC of 0.66, AUROC of 0.736, MCC of 0.36, and F1-score of 0.658 (Table 4B). The combined ROC curves for internal and external validation experiments are shown in Figure 4B-C.

**Table 4A:** Top internal validation results using derivation set (GSE25066 – 170 patients)

|  | Sensitivity<br>(95% CI) | Specificity<br>(95% CI) | AUROC<br>(95% CI) | MCC<br>(95% CI) |
| --- | --- | --- | --- | --- |
| ROS_ET_30 | 0.768 (0.733-0.8) | 0.916 (0.878-0.94) | 0.771 (0.733-0.82) | 0.549 (0.515-0.6) |
| ROS_ET_25 | 0.712 (0.711-0.789) | 0.884 (0.81-0.891) | 0.753 (0.735-0.782) | 0.525 (0.499-0.566) |
| SMOTE_ET_25 | 0.782 (0.732-0.802) | 0.916 (0.87-0.94) | 0.772 (0.744-0.809) | 0.55 (0.49-0.61) |
| SMOTE_ET_20 | 0.743 (0.697-0.788) | 0.833 (0.778-0.867) | 0.771 (0.75-0.793) | 0.533 (0.491-0.576) |
| ADASYN_ET_30 | 0.772 (0.705-0.81) | 0.84 (0.79-0.89) | 0.769 (0.744-0.8) | 0.547 (0.533-0.58) |
| SMOTE_KNN_30 | 0.728 (0.699-0.755) | 0.7 (0.65-0.74) | 0.737 (0.71-0.756) | 0.511 (0.46-0.529) |
| Tomek_KNN_29 | 0.766 (0.723-0.8) | 0.71 (0.67-0.75) | 0.755 (0.712-0.8) | 0.52 (0.487-0.545) |
| CC_KNN_29 | 0.743 (0.723-0.767) | 0.76 (0.72-0.82) | 0.744 (0.715-0.756) | 0.5 (0.44-0.53) |

ROS: Random OverSampling; ET: ExtraTress; RF: Random Forests; KNN:K-nearest neighbors; Tomek: TomekLinks; CC: Cluster Centroids

**Table 4B:** Top external validation results on microarray and RNA-seq datasets – Combined [GSE20271, GSE20194, GSE31519 (194 patients)] and GSE106977 (119 patients)

| **GSE20271, GSE20194, GSE31519 (Microarray - 194 patients)** |  |  |  |  |  |  |  |
| --- | --- | --- | --- | --- | --- | --- | --- |
| Classifier-sampling technique-number of features | Sensitivity | Specificity | AURPC | Balanced Accuracy | AUROC | MCC | F1 |
| ROS_ET_30 | 0.908 | 0.527 | 0.744 | 0.718 | 0.772 | 0.483 | 0.751 |
| ROS_ET_25 | 0.925 | 0.541 | 0.754 | 0.733 | 0.777 | 0.52 | 0.766 |
| SMOTE_ET_25 | 0.925 | 0.514 | 0.745 | 0.719 | 0.8 | 0.497 | 0.754 |
| SMOTE_ET_20 | 0.908 | 0.581 | 0.764 | 0.745 | 0.799 | 0.53 | 0.775 |
| ADASYN_ET_30 | 0.917 | 0.541 | 0.752 | 0.729 | 0.798 | 0.508 | 0.762 |
| **GSE106977 (Microarray - 119 patients)** |  |  |  |  |  |  |  |
| ROS_ET_30 | 0.534 | 0.739 | 0.694 | 0.637 | 0.64 | 0.269 | 0.616 |
| ROS_ET_25 | 0.534 | 0.696 | 0.679 | 0.615 | 0.658 | 0.225 | 0.601 |
| SMOTE_ET_25 | 0.562 | 0.674 | 0.68 | 0.618 | 0.641 | 0.23 | 0.61 |
| SMOTE_ET_20 | 0.425 | 0.783 | 0.674 | 0.604 | 0.606 | 0.212 | 0.558 |
| ADASYN_ET_30 | 0.411 | 0.783 | 0.67 | 0.597 | 0.658 | 0.2 | 0.548 |
| **GSE163882 (RNA seq - 90 patients)** |  |  |  |  |  |  |  |
| Tomek_KNN_29 | 0.923 | 0.368 | 0.66 | 0.646 | 0.736 | 0.36 | 0.658 |
| SMOTE_KNN_30 | 0.577 | 0.711 | 0.667 | 0.644 | 0.722 | 0.285 | 0.635 |
| ROS_KNN_29 | 0.462 | 0.816 | 0.668 | 0.639 | 0.699 | 0.288 | 0.604 |
| CC_KNN_29 | 0.827 | 0.421 | 0.647 | 0.624 | 0.711 | 0.273 | 0.639 |
| CC_KNN_30 | 0.577 | 0.684 | 0.657 | 0.631 | 0.678 | 0.259 | 0.624 |

### Survival analysis of the selected marker genes displays a correlation between higher expression and better survival

After the derivation and validation analysis, we analyzed all top 30 features to test their association with RFS using the online KMplotter tool. Kaplan-Meier RFS analysis identified better survival with high expression of the gene markers and vice versa. (Figure 5A; log-rank P value = 0.011; HR: 0.52). Additionally, the top 30 TIL-based markers established through our rigorous feature selection pipeline showed higher expression in the chemosensitive group of cells than the chemoresistant ones (Figure 5B; Wilcoxon test P value < 0.05). As established through our enrichment analysis, the marker genes were associated with immune cell activation and proliferation. In chemosensitive patients, where the immune system is better able to support the effects of chemotherapy, these genes were more highly expressed. These observations align with previous studies that reported an association between overexpression of immunological gene signatures and better RFS in early-stage chemotherapy-treated TNBC patients (2,52).

**Figure 5:**
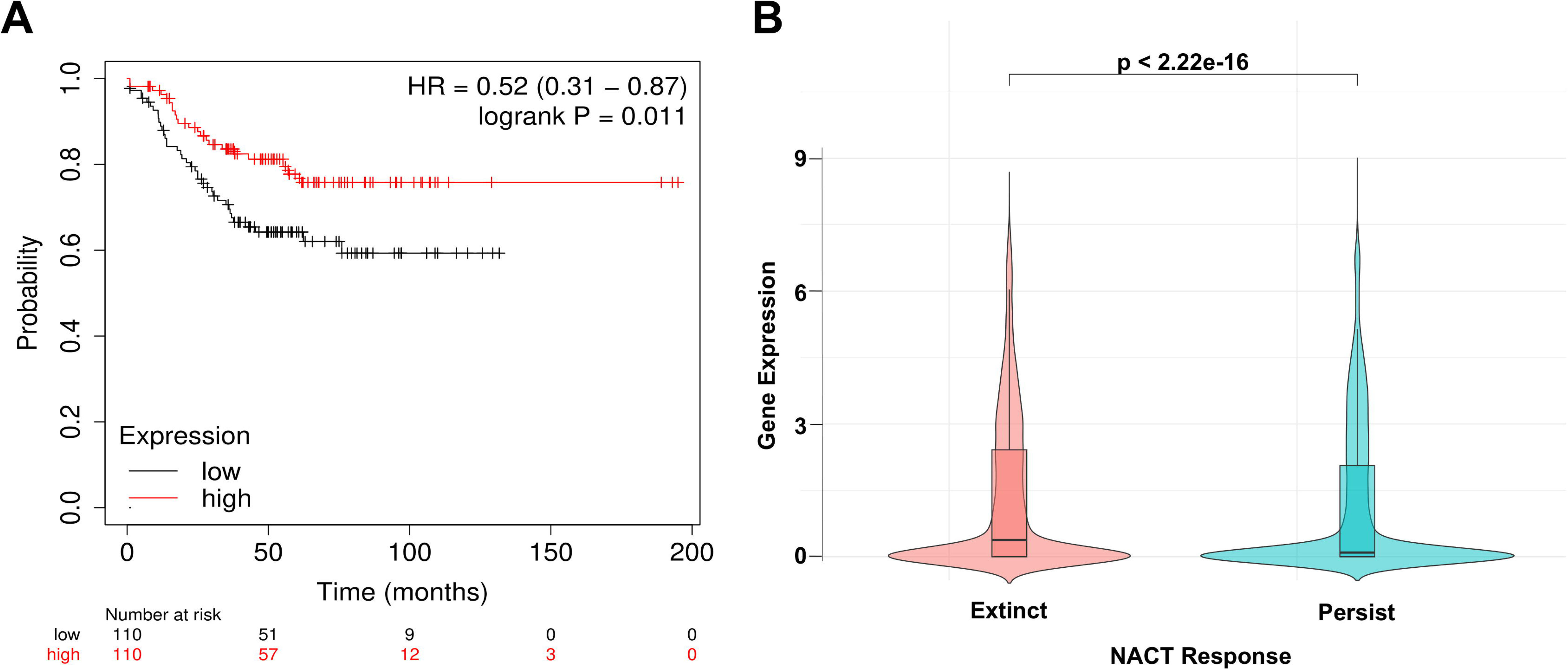
Survival analysis using the top-ranked markers validated using different internal and external cohorts. (A) Kaplan-Meier RFS analysis identified better survival with high expression of the gene markers (B) Dependence between median expression and chemotherapy response using the top 30 genes identified using our technique.

## Discussion

Triple-negative breast cancer (TNBC) is a heterogeneous disease with an immune microenvironment that affects chemotherapy response in patients undergoing neoadjuvant chemotherapy (NAC). Tumor-infiltrating lymphocytes (TILs) play a key role in modulating this microenvironment. This study analyzed single-cell and bulk RNA sequencing from public datasets to identify NAC response markers within the TIL subpopulation. Based on gene expression data, we used machine learning models to predict pathological complete response (pCR) versus residual disease (RD) in TNBC patients. Our extensively validated models showed that higher expression of these markers was associated with better survival outcomes in NAC-treated patients.

Initial enrichment analysis of single-cell data revealed dysregulation in immune function, cell adhesion, and extracellular matrix (ECM) organization, aligning with previous studies (53–55). Suppressed immune infiltration is known to predict therapy response and survival outcomes (54,55). Similarly, alterations in tumor-derived ECM, biochemically distinct from normal ECM, can increase hypoxia and metabolic stress, impairing therapy response (53,56). Neutrophil degranulation, a significantly altered REACTOME pathway in our analysis, has been shown to promote immunosuppressive activity in TNBC patients, particularly via myeloid-derived suppressor cells (MDSCs) (57). Conversely, enriched gene expression in the TIL subpopulation indicated heightened lymphocyte activity and immune suppression, characteristic of the TNBC transcriptome (58).

Machine learning algorithms, designed to identify complex patterns in gene expression, have emerged as a leading approach for biomarker discovery (59–61). However, gene expression data often faces the challenge of dimensionality. For example, our microarray derivation dataset includes only 170 samples but over 14,000 features. To address this, we used a “result set filtering” strategy from Perscheid et al. (62), which incorporates prior biological knowledge to focus on biologically relevant genes and reduce the feature search space. By intersecting differentially expressed genes from TIL subpopulations with a curated list of genes linked to chemoresistance in NAC-treated breast cancer patients, we identified 226 genes, mainly involved in cellular division pathways. As cancer is marked by cell cycle dysregulation (63), which drives tumor growth, intratumoral TILs are associated with heightened immune response and cell proliferation (64). Notably, high expression of cell cycle regulators like CENPF (one of the 226 genes) correlates with poor chemotherapy outcomes (65).

Five key genes—FOXM1, CCNB1, CCNA2, BIRC5, and TOP2A—were identified through consensus hub gene identification techniques, all linked to poor TNBC prognosis and survival ((66–69), Figure 4D). For example, CCNB1, a cyclin family member, plays a critical role in cancer progression and is an independent prognostic factor in node-negative breast cancer (70–72). Likewise, CCNA2, a cell cycle regulator, is associated with E2F transcription factor activity, driving TNBC proliferation (73). Notably, BIRC5, expressed in 45-90% of TNBC patients, belongs to the inhibitor of apoptosis protein (IAP) family, which regulates cell death and proliferation. It is a key prognostic marker for chemoresistance in TNBC (69).

We employed a recursive feature elimination approach to select genes implicated in chemoresistant TNBC samples, as described in the methods section. Several of the 30 genes identified as prognostic markers for NAC response have established roles in TNBC. Furthermore, high expression of these marker genes, observed mainly in the chemosensitive group, is also associated with improved RFS. Higher expression in chemosensitive patients could reflect a generally more favorable tumor microenvironment rather than directly predicting chemotherapy response attesting to the prognostic relevance of the markers. Among the top-ranked genes, Nuclear Factor I B (NFIB) promotes cell proliferation and therapy resistance in triple-negative and ER-negative breast cancers (74,75). The Vascular Endothelial Growth Factor (VEGF) family, particularly VEGFA, is a key regulator of angiogenesis, with high expression linked to poor prognosis and lower 5-year survival rates in 60% of TNBC cases (76). The S100 calcium-binding protein family, including S100B and S100P, contributes to the pro-inflammatory tumor microenvironment (TME) and serves as independent prognostic markers for metastasis and therapy resistance in TNBC (77,78). Aberrant expression of DEPDC1, NPY1R, STAT3, and GSR is associated with cancer cell invasion, proliferation, immunosuppression, cisplatin detoxification, chemoresistance, metastasis, and activation of the PI3K/AKT/mTOR pathway, presenting them as novel therapeutic targets (79–81). In silico analysis of the breast cancer transcriptome revealed that CRIM1 expression is lowest in TNBC among all breast cancer subtypes and regulates several EMT-related factors (82). Similarly, public dataset analysis identified the SFRP1 gene, strongly associated with the TNBC subtype, as a potential marker for predicting chemotherapy response (83). In a recent study, PTPRC has been implicated in paclitaxel resistance in TNBC cell lines by modulating CD8+ T cell infiltration (84).

Our study identified key prognostic factors and immunotherapy targets for TNBC, including CDKN2A (85), IGFBP4 (86), CLDN3 (87), RPL31 (88), and CXCL11 (89), which may guide future therapeutic strategies. The transcription factor E2F3, crucial for epithelial-to-mesenchymal transition in breast cancer, suppresses tumor growth and metastasis when its expression is reduced (90). Conversely, elevated DEK expression is linked to cell differentiation, metastasis, poor survival, and EMT promotion in TNBC (89). Additionally, abnormal SAA1 expression drives adipocyte reprogramming, leading to adipocyte infiltration, stemness, and inflammation in TNBC patients (91).

Among the top-ranked features, increased expression of the tumor suppressor gene Nucleolar protein 7 (NOL7) is linked to poor prognosis and overall survival in several cancers, including TNBC (34,92,93). While Histone Acetyltransferase 1 (HAT1) has not been studied in TNBC, it regulates cell proliferation, cancer immunity, and transcription of PD-L1 in pancreatic cancer, serving as a predictive marker for immune checkpoint therapy (94). Similarly, although not directly implicated in TNBC, downregulation of SEC13 has been associated with aberrant TGF-B signaling in gastric cancer, indicating drug resistance (95). The Trafficking From ER to the Golgi Regulator (TFG) gene, involved in protein distribution, apoptosis, and cell growth, is a therapeutic target in cancers such as secretory breast carcinoma (96). Upregulated expression of Chaperonin containing TCP1 subunit 5 (CCT5) in p53-mutated breast cancers may confer resistance to docetaxel and identify patients unlikely to respond to therapy (97). Elevated levels of transmembrane proteins such as GNG12 and TMEM14B contribute to cancer progression, including TNBC, by driving cell proliferation, inflammatory and immune responses, and key signaling pathways (98,99).

### Limitations

Our study has some limitations: first, the small sample size and the need for consistent clinical information, such as age, tumor grade, stage, etc., across all derivation and validation datasets prevented us from building an integrated model including both gene and clinical information. Second, due to the lack of single-cell datasets with chemotherapy response labels from TNBC patients, we used microarray and RNA-seq data to derive and validate our interpretations of the marker genes. Third, we used a novel hybrid method of selecting the most informative features from a single-cell population using microarray data. Other conventional feature selection methods must be explored to establish their validity. Nonetheless, we validate our marker set on a combined sample size of approximately 400 patients across various sequencing platforms and technologies (microarray: GPL570, GPL96, GPL17586, and RNA-seq: GPL18573) to reduce bias in the model building and interpretation process.

## Conclusion

The present study introduces an *in silico* method for identifying key prognostic markers associated with NAC response in pre-treated TNBC patients. This method analyzes TIL subpopulations using single-cell expression data and validates the findings using bulk RNA-seq datasets. The list of 30 features was derived from a rigorous feature selection, derivation, and validation pipeline wherein we measured the variability of the selected genes to ensure a stable set of generalizable features.

Most derived genes are established markers with varying functionalities and prognostic relevance to NAC response among TNBC patients. The resulting derivation AUROC of 0.78 and validation AUROCs of 0.8, 0.658, and 0.736 across five independent test sets spanning different platforms and sequencing technologies indicate the prognostic significance of these markers in reliably predicting the response to chemotherapy among NAC-administered TNBC patients. Furthermore, these genes serve as markers of a favorable immune microenvironment rather than being directly responsible for chemoresistance. While further in silico validation on closely related datasets and experimental studies would be desirable to confirm our finding, our 30-gene classifier is a promising tool for stratifying TNBC patients at risk of developing chemoresistance to NAC. Our work paves the way for personalized treatment strategies and improved clinical outcomes, with a key role for the marker identified herein as potential targets for therapeutic interventions.

## Acknowledgements

SB acknowledges funding support for the research of this work from the Centre for Integrative Biology and Systems mEdicine (IBSE), Indian Institute of Technology Madras.

---

**Supplementary Figure (1A-C):**
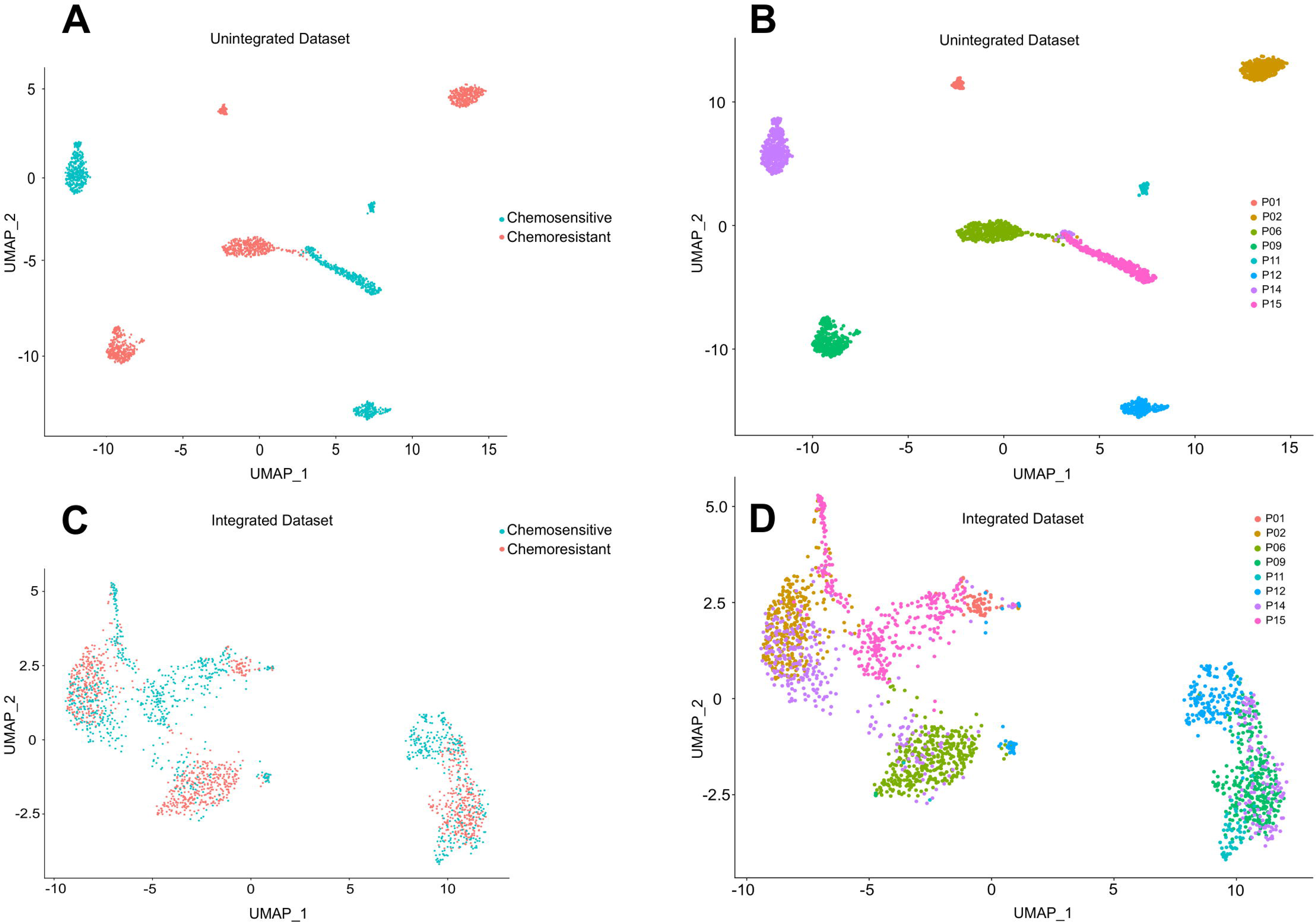
UMAP showing the effect of integration on our single-cell dataset using NAC response as labels. (1B-D) UMAP shows the effect of integration on our single-cell dataset using patient identities as labels.

**Supplementary Figure 2A:**
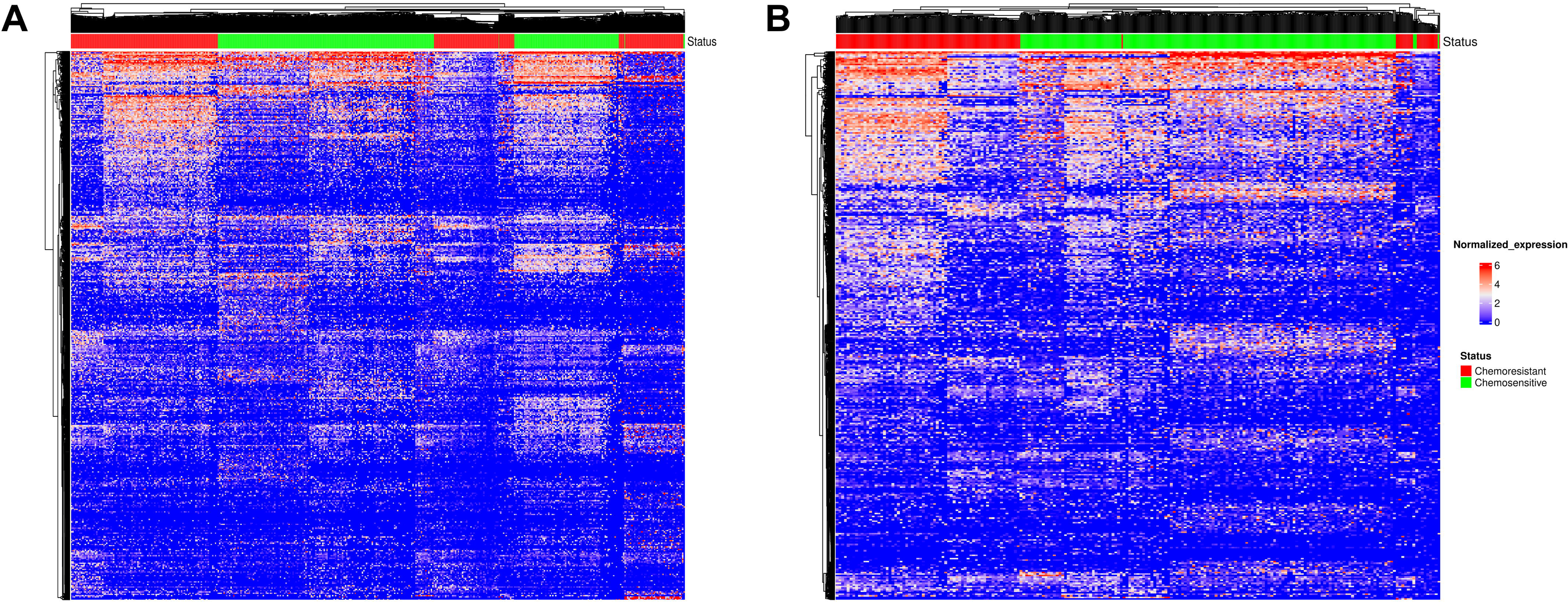
Heatmap showing the expression values of the differentially expressed genes from the original set of 2640 cells cluster based on NAC response labels. (2B): Heatmap showing the expression values of the differentially expressed genes from the set of 860 TIL subpopulations of cells cluster based on NAC response labels.

